# Non-canonical HIF-1 stabilization is essential for intestinal tumorigenesis

**DOI:** 10.1101/273045

**Authors:** Nadine Rohwer, Sandra Jumpertz, Merve Erdem, Antje Egners, Klaudia T. Warzecha, Athanassios Fragoulis, Anja A. Kühl, Rafael Kramann, Sabine Neuss, Ines Rudolph, Tobias Endermann, Christin Zasada, Ivayla Apostolova, Marco Gerling, Stefan Kempa, Russell Hughes, Claire E. Lewis, Winfried Brenner, Maciej B. Malinowski, Martin Stockmann, Lutz Schomburg, William Faller, Owen Sansom, Frank Tacke, Markus Morkel, Thorsten Cramer

## Abstract

The hypoxia-inducible transcription factor HIF-1 is appreciated as a promising target for cancer therapy. However, conditional deletion of HIF-1 and HIF-1 target genes in cells of the tumor microenvironment can result in accelerated tumor growth, calling for a detailed characterization of the cellular context to fully comprehend HIF-1’s role in tumorigenesis. We dissected cell type-specific functions of HIF-1 for intestinal tumorigenesis by lineage-restricted deletion of the *Hif1a* locus. Intestinal epithelial cell-specific *Hif1a* loss reduced activation of wnt/β-catenin, tumor-specific metabolism and inflammation, significantly inhibiting tumor growth. Deletion of *Hif1a* in myeloid cells reduced the expression of fibroblast-activating factors in tumor-associated macrophages resulting in decreased abundance of tumor-associated fibroblasts and robustly reduced tumor formation. Interestingly, hypoxia was detectable only sparsely and without spatial association with nuclear HIF-1α in intestinal adenomas, pointing towards a functional importance of hypoxia-independent, i.e. non-canonical HIF-1 stabilization that has not been previously appreciated. This adds a further layer of complexity to the regulation of HIF-1α and suggests that hypoxia and HIF-1α stabilization can be uncoupled in cancer. Collectively, our data show that HIF-1 is a pivotal pro-tumorigenic factor for intestinal tumor formation, controlling key oncogenic programs in both the epithelial tumor compartment and the tumor microenvironment.

## Introduction

Colorectal cancer (CRC) ranks as the third most commonly diagnosed tumor in both sexes in the western world, causing a substantial amount of cancer-associated morbidity and mortality (1). In the last decade, molecular-targeted drugs have -in combination with conventional therapies-resulted in improved outcome of CRC patients (2). However, as also noted frequently for other solid tumors, these new generation treatments rarely result in improved outcome, but rather benefit small subgroups of CRC patients (3). Taken together, an urgent need to identify novel therapy targets exists, demonstrating the necessity for detailed characterization of the molecular CRC pathogenesis.

The hypoxia-inducible transcription factor 1 (HIF-1) is positively associated with the malignant progression of various tumor entities (4-7). While the role of HIF-1 for the pathogenesis of CRC has been addressed by different groups, results are conflicting and the precise function of HIF-1 for intestinal tumorigenesis remains elusive. Stabilization of HIF-1α, the regulatory component of HIF-1, in human CRC has been reported via immunohistochemistry, suggesting a pro-tumorigenic function (8-10). In line with these findings, inhibition of HIF-1α in human CRC cell lines resulted in reduced xenograft growth (11, 12). Furthermore, pharmaceutical inhibition of HIF-1α led to diminished growth of autochtonous as well as allograft murine CRC models via reduced angiogenesis and macrophage infiltration (13). On the other hand, transgenic activation of HIF-1 in intestinal epithelial cells (IEC) remained without effect on tumor formation in murine models of sporadic and colitis-associated colon cancer (14, 15). Moreover, IEC-specific loss of *Hif1a* did not affect tumor frequency in a chemical model of proximal intestinal tumorigenesis (16). These results illustrate that the precise role of HIF-1 for intestinal tumorigenesis remains elusive thus far.

While HIF-1 is widely appreciated as a promising target for cancer therapy (17), conditional deletion of HIF-1 and HIF-1 target genes in cells of the tumor microenvironment can result in accelerated tumor growth (18-20). Against this background, a comprehensive analysis of HIF-1’s role in tumorigenesis in order to translate the findings into the clinic can only be achieved by considering the cell- and tissue-specific context. Here, we present a detailed deconstruction of HIF-1’s role in CRC initiation and progression in IEC- and myeloid cell-specific *Hif1a* knock-out mice (21, 22) using two murine tumor models: Chemically-induced colon tumors (AOM/DSS (combination of <u>a</u>z<u>o</u>xy<u>m</u>ethane plus<u>d</u>extran <u>s</u>ulfate <u>s</u>odium for colitis-associated carcinogenesis) (23)) and the genetic APC^min^ model (24). This experimental approach enabled us to address the role of HIF-1 in neoplastic epithelial cells as well as in innate immune cells of the tumor microenvironment. We were able to unravel numerous specialized functions of HIF-1. In IECs, HIF-1 serves to induce inflammation, control wnt/β-catenin activity and regulate tumor-specific metabolism. In addition, we identified myeloid HIF-1 as essential for the activation of tumor-associated fibroblasts. Of note, intratumoral hypoxia was a rare event, pointing towards an importance of hypoxia-independent, i.e. non-canonical HIF-1 stabilization for intestinal tumorigenesis (25). Collectively, our data show that HIF-1 is a pivotal pro-tumorigenic factor, controlling key oncogenic programs in both the epithelial tumor compartment and the tumor microenvironment.

## Results

### Epithelial *Hif1a* controls intestinal tumor growth on multiple levels

IEC-specific loss of *Hif1a* (termed Hif1a^IEC^) was established by breeding *villin-cre* mice (26) with animals harbouring floxed *Hif1a* alleles (22). Conditional *Hif1a* deletion was characterized on various levels and found to be highly efficient (Figure S1A). Tumor size in both the AOM/DSS and APC^min^ model was significantly decreased in Hif1a^IEC^ mice (Figure 1A). We sought to identify the underlying molecular mechanisms and first characterized the activity of the wnt/β-catenin pathway, the central oncogenic driver of CRC. As can be seen in figure 1B-D, wnt-dependent gene expression is significantly reduced in adenomas of Hif1a^IEC^ mice in both tumor models, suggesting a functional role of HIF-1 for wnt/β-catenin activity in this setting. Next, we addressed the role of *Hif1a* for tumor-specific glucose metabolism as metabolic reprogramming represents an emerging hallmark of cancer and several glycolytic enzymes are transcriptionally controlled by HIF-1 (27). Mass spectrometry-based analysis of intratumoral glucose levels, routing of ^13^C-labelled glucose as well as *in vivo* imaging by PET/CT revealed diminished uptake and metabolization into lactate of glucose in tumors of Hif1a^IEC^ mice (Figure 1E, F). Finally, we performed a comprehensive analysis of inflammatory activity as chronic inflammation is pro-tumorigenic in CRC (28). Hif1a^IEC^ mice displayed reduced loss of body weight (Figure 2A) and colitis activity after DSS challenge (Figure 2B). Intestinal gene expression and protein secretion of pivotal pro-inflammatory factors were significantly lower in DSS-treated Hif1a^IEC^ mice (Figure 2C, D). Experiments with small intestinal organoids were conducted to further address the contribution of IECs in this setting. After assuring efficient *Hif1a* deletion (Figure S2), organoids were subjected to different pro-inflammatory stimuli. These experiments nicely confirmed that the pro-inflammatory response of IECs critically depends on *Hif1a* (Figure 2E). Taken together, these data point towards a complex role for *Hif1a* in IECs during intestinal tumor formation, comprising inflammation, wnt/β-catenin activity and metabolic reprogramming.

**Figure 1.**
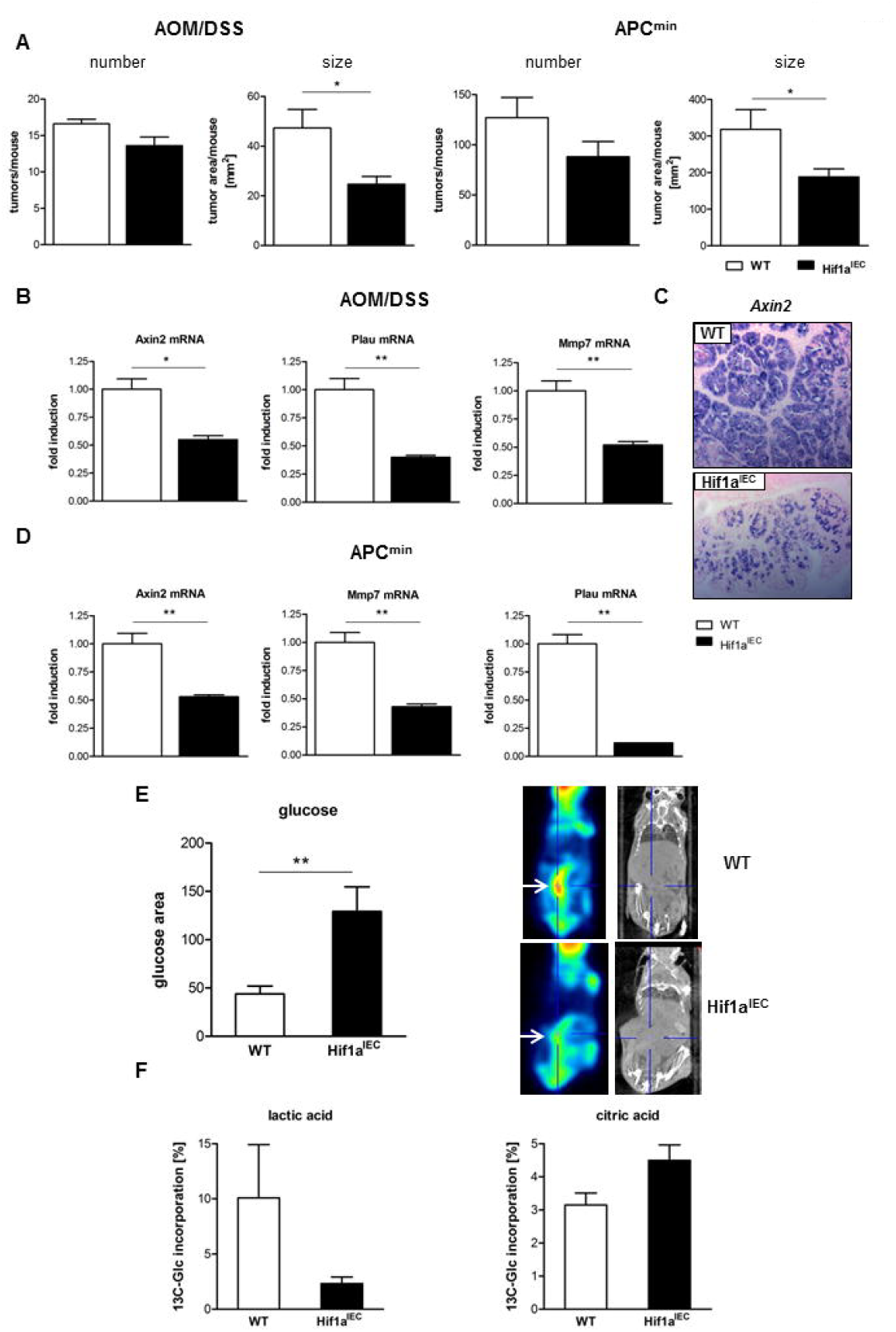
HIF-1α in IECs controls intestinal tumor formation, wnt/β-catenin activity and glucose metabolism. (**A**) Tumor number and size of WT and Hif1a^IEC^ mice in the AOM/DSS (left, *n* = 4-6 per group) and APC^min^ model (right, *n* = 7 per group). (**B**) Relative mRNA expression of wnt/β-catenin target genes in colon tumors of WT and Hif1a^IEC^ mice following AOM/DSS treatment, *n* = 3 per group. (**C**) RNA *in-situ* hybridization of the wnt/β-catenin target gene *Axin2* in AOM/DSS adenomas of WT and Hif1a^IEC^ mice. (**D**) Relative mRNA expression of wnt/β-catenin target genes in WT and Hif1a^IEC^ APC^min^ tumors, *n* = 3 per group. (**E**) Glucose levels (left) and FDG-PET/CT analysis (right) of AOM/DSS-induced colon tumors (white arrow) of WT (*n* = 5) and Hif1a^IEC^ (*n* = 8) mice. (**F**) Metabolization of glucose into lactate (left, *n* = 3-5 per group) and citrate (right, *n* = 13 per group) in colon tumors of WT and Hif1a^IEC^ mice following AOM/DSS treatment determined by stable isotope-resolved metabolomics. Data are represented as mean + SEM. *p<0.05; **p<0.01 by unpaired Student’s t test.

**Figure 2.**
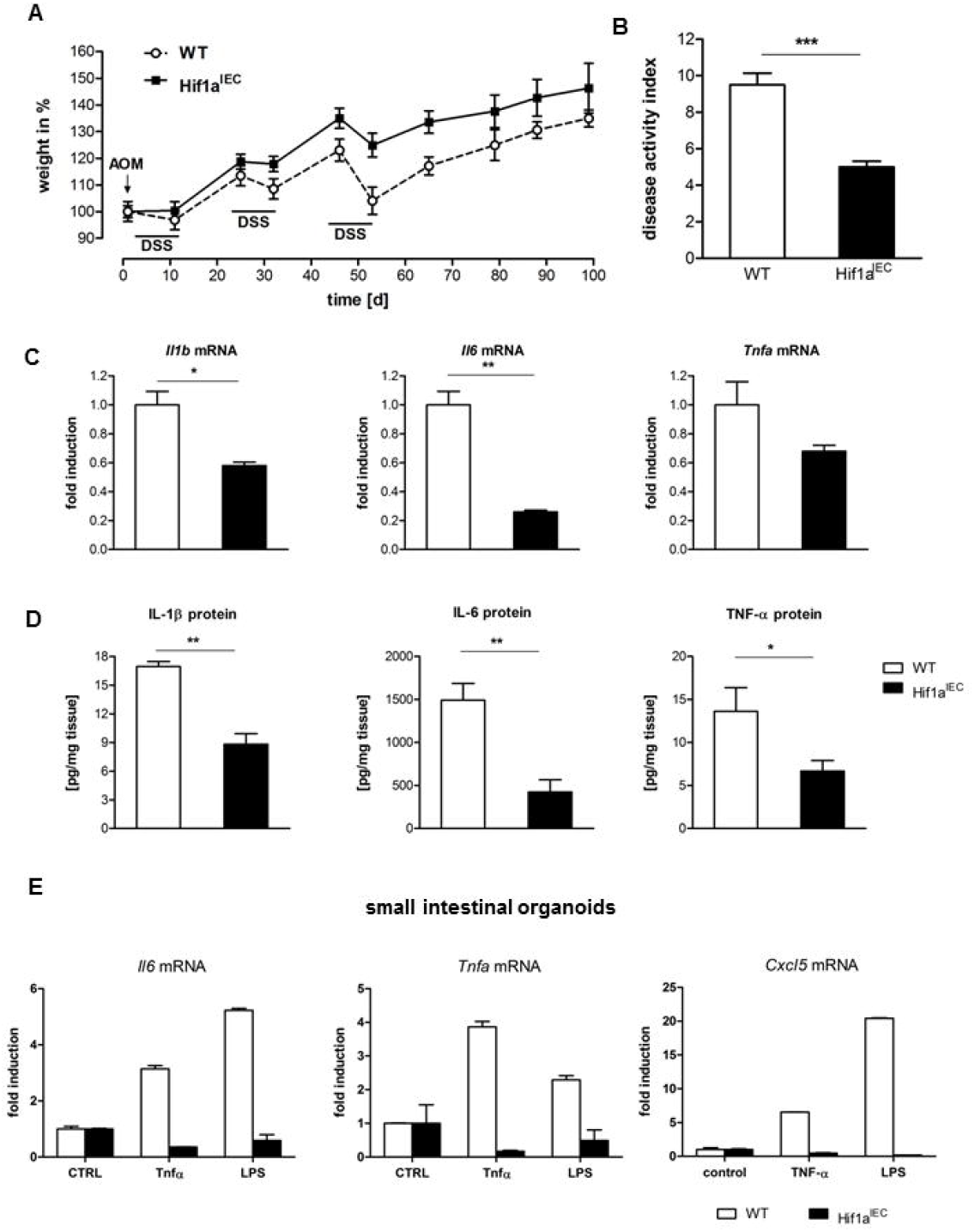
IEC-specific deletion of *Hif1a* reduces intestinal inflammation. (**A**) Body weight change of WT and Hif1a^IEC^ mice following AOM/DSS treatment (*n* = 14 per group), (**B**) disease activity index (*n* = 4-5 per group), (**C**) relative mRNA expression of inflammatory genes in colon tissues (*n* = 3 per group) and (**D**) secretion of cytokines by colonic explants of WT and Hif1a^IEC^ mice on day 6 of acute DSS treatment (*n* = 4-5 per group). (**E**) Pro-inflammatory cytokine mRNA expression in small intestinal organoids from WT and Hif1a^IEC^ mice (*n* = 3 per group). Shown are mean and standard deviation of technical duplicates from one representative experiment. Unless indicated otherwise, data are represented as mean + SEM. *p<0.05; **p<0.01; ***p<0.001 by unpaired Student’s t test.

### *Hif1a* in myeloid cells controls intestinal tumor formation without affecting inflammation

Macrophages in the tumor microenvironment exert a number of tumor-supporting functions and are positively associated with the malignant phenotype (29). We and others have shown that *Hif1a* is of pivotal importance for various aspects of macrophage function (21, 30). Against this background, we sought to characterize the role of *Hif1a* for intestinal tumor formation in macrophages. We found that myeloid cell-specific loss of *Hif1a* (termed Hif1a^MC^) resulted in a highly significant reduction of both tumor number and size (Fig. 3A). In order to identify the underlying mechanisms, we first analyzed the inflammatory response. Rather unexpectedly, various assays of inflammation (e.g. determination of weight loss (Fig. 3B), disease activity index (Fig. 3C)) as well as gene expression and protein production of established pro-inflammatory markers (Fig. 3D, E, F)) failed to show a difference of inflammatory activity between wildtype and Hif1a^MC^ mice.

**Figure 3.**
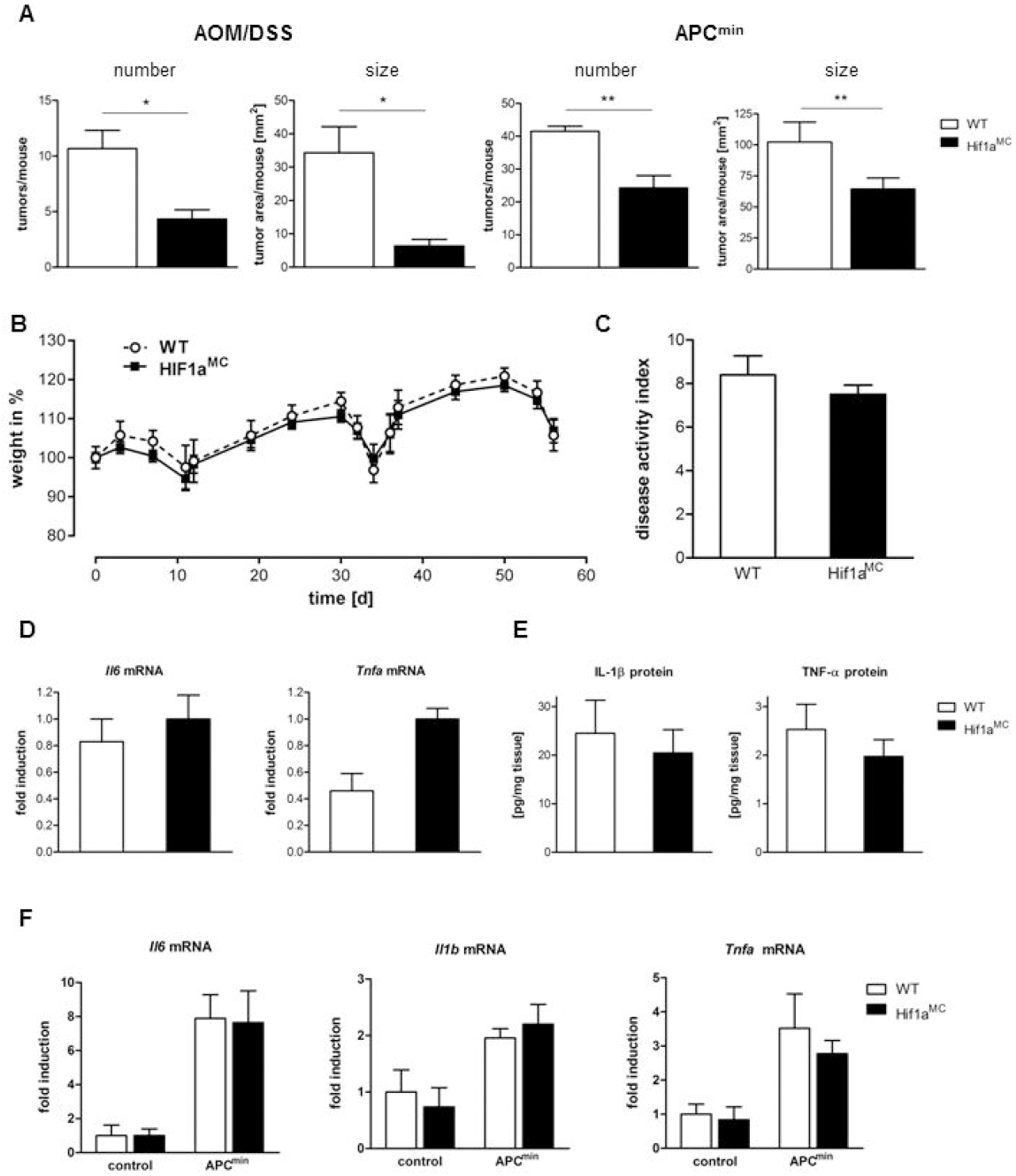
*Hif1a* in myeloid cells controls intestinal tumor formation without affecting inflammation. (**A**) Tumor number and size in WT and Hif1a^MC^ mice after AOM/DSS administration (left, *n* = 3 per group) and in the APC^Min^ model (right, *n* = 4 per group). (**B**) Body weight change of WT (*n* = 7) and Hif1a^MC^ (*n* = 8) mice following AOM/DSS treatment. (**C**) Disease activity index (*n* = 5-6 per group), (**D**) Relative mRNA expression of inflammatory genes in colon tissues (*n* = 3 per group) and (**E**) Secretion of cytokines by colonic explants of WT and Hif1a^MC^ mice on day 6 of acute DSS treatment (*n* = 5 per group). (**F**) Relative mRNA expression of inflammatory genes in intestinal tissues of WT and Hif1a^MC^ control (*n* = 3-4 per group) and APC^min^ (n = 7-8 per group) mice. Data are represented as mean + SEM. *p<0.05; **p<0.01 by unpaired Student’s t test.

### *Hif1a*-deficient tumor-associated macrophages migrate and function normally

Next, we determined intratumoral macrophage numbers via immunohistochemistry (IHC). While macrophage abundance was clearly greater in adenomas compared to surrounding normal mucosa (not shown), no difference was detectable between the genotypes (Figure 4A). Intestinal leukocyte subsets were subsequently analyzed in more detail by flow cytometry. Tumor-bearing mice displayed a prominent increase in CD11c+ macrophages and CD11b+ dendritic cells, while CD11c-macrophages were reduced. The changes in myeloid subsets were comparable between wildtype and Hif1a^MC^ mice, except for a slightly more pronounced reduction of CD11c-intestinal macrophages in tumor-bearing Hif1a^MC^ mice (Figure 4B). Intestinal lymphoid cell populations did not differ significantly between wildtype and Hif1a^MC^ mice (Figure 4B). Notably, tumor-bearing animals showed typical alterations of extraintestinal myeloid cells (31), including increase of monocytes in blood and bone marrow and accumulation of Gr1+ CD11b+ myeloid-derived suppressor cells (MDSC) in bone marrow, but not spleen (Figure S3). Extraintestinal myeloid cells did not differ between wildtype and Hif1a^MC^ mice. Next, we decided to analyze the direct tumor-supporting action of macrophages as these cells secrete numerous pro-tumorigenic factors (32). To this end, spheroids of APC^min^ adenomas were stimulated with conditioned medium (CM) from primary murine macrophages. While we were able to detect a significant growth-promoting effect of macrophage CM, the loss of *Hif1a* did not affect the outcome (Figure 4C). As macrophages represent pivotal modulators of stem-like/progenitor cells (33), which are critical drivers of colonic carcinogenesis (34), we decided to quantify these cells in our experimental setting. Visualization of *Lgr5* and *Prox1*, two established progenitor/stem-cell markers in intestinal tumors (35, 36), demonstrated robust presence of these cells in AOM/DSS adenomas, albeit without differences between the genotypes (Figure S4A). Furthermore, we took into consideration that other cells of myeloid origin might underlie the reduced tumor formation in Hif1a^MC^ mice. While mast cells can influence intestinal tumorigenesis under certain experimental conditions (37), no difference in intratumoral abundance of this cell type was noted, arguing against a functional importance of mast cells in our setting (Figure S4B).

**Figure 4.**
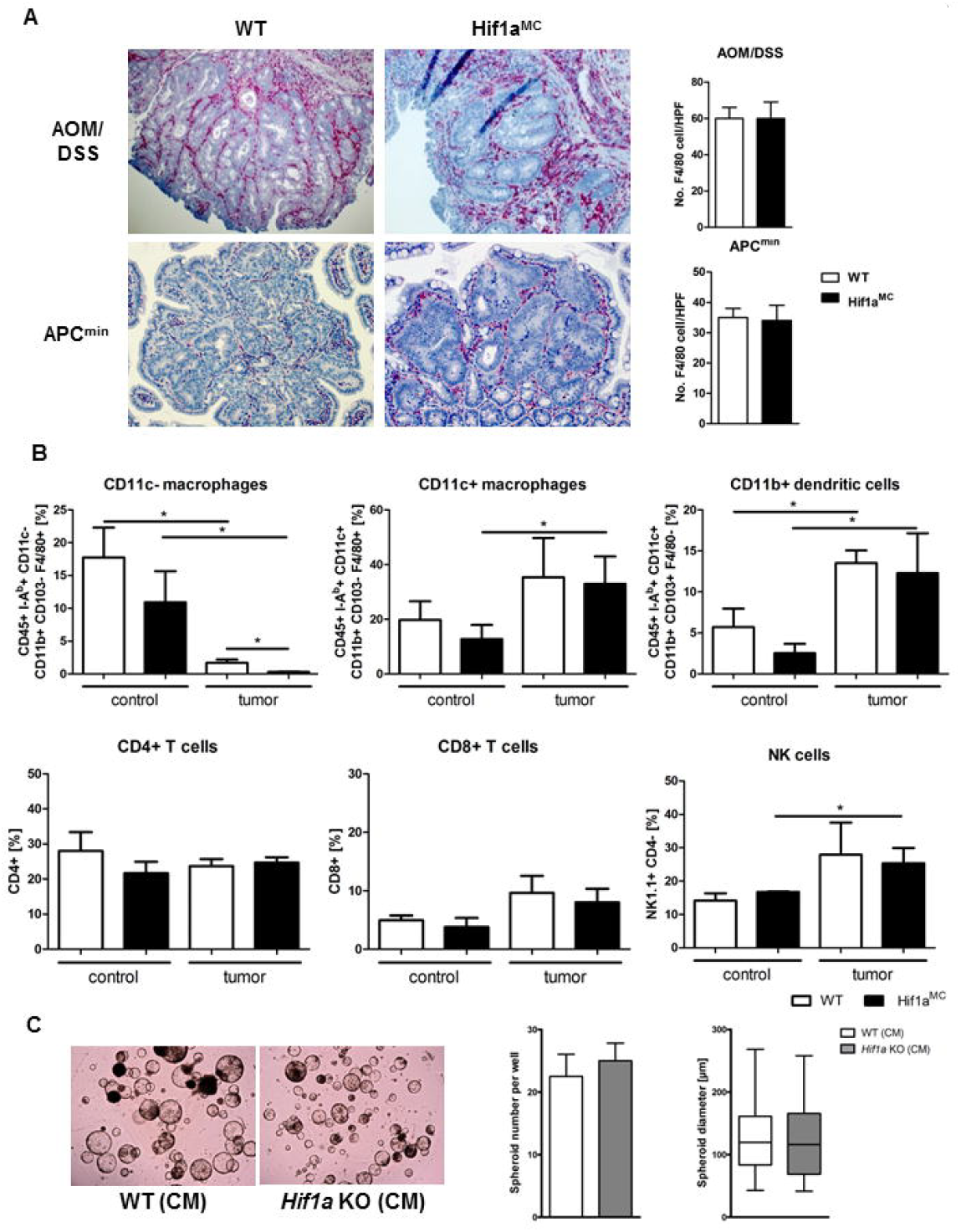
Tumoral abundance of macrophages and stimulation of adenoma growth *ex vivo* is not affected by loss of *Hif1a.* (**A**) Representative F4/80 stainings of intestinal sections obtained from AOM/DSS-treated (above) and APC^min^ (below) WT and Hif1a^MC^ mice. Right, quantification of F4/80-positive cells (*n* = 6 per group). (**B**) Total leukocytes were isolated from small intestines of wildtype and Hif1a^MC^ mice remaining either untreated or bearing APC^min^ adenomas and analyzed by flow cytometry (*n* = 4 per group). Relative numbers of different subtypes are shown. Flow cytometric analyses of the CD11c-macrophage subset (I-Ab+, CD11c-, CD11b+, CD103-, F4/80+), the CD11c+ macrophage subset (I-Ab+, CD11c+, CD11b+, CD103-, F4/80+) and the CD11b+ dendritic cell subset (I-Ab+, CD11c+, CD11b+, CD103+, F4/80-) were performed. Lymphoid leukocytes of the small intestine were subdivided into T cells (CD4+, CD8+) and NK cells (CD4-, NK1.1+). (**C**) Effect of macrophage conditioned media (CM) on spheroid formation from APC^min^ adenomas. Left, representative image of spheroids after stimulation. Quantification of spheroid number (middle) and diameter (right) after stimulation with conditioned media from WT and *Hif1a*-KO macrophages (*n* = 3 per group). Data in **A** and **B** are represented as mean + SEM. *p<0.05 by unpaired Student’s t test. **C** right shows mean ± SD.

### *Hif1a* in myeloid cells is essential for the activation of tumor-associated fibroblasts

Myeloid cells in the tumor microenvironment are known to interact with various cell types, including tumor-associated fibroblasts (TAF). These cells are abundant in human CRC and significantly impact on disease progression (38, 39). TAFs were readily detectable via IHC in the stroma of APC^min^ and AOM/DSS-induced adenomas (Figure 5A). Strikingly, deletion of *Hif1a* in myeloid cells resulted in greatly reduced numbers of TAFs in both tumor models (Figure 5A). This result pointed towards a crucial role of *Hif1a* in myeloid cells for TAF development. Interestingly, the importance of macrophages for fibroblast activation in the context of wound healing is well established (40, 41). To analyze this further, we focussed on alternatively activated macrophages (AAM) given their central role in the control of organ fibrosis (40). Expression of various pro-fibrotic genes in AAMs was readily detectable, but loss of *Hif1a* remained without effect (Figure S5A). To achieve experimental data that more closely resemble the *in vivo* situation, we analyzed pro-fibrotic factor expression in tumor-associated macrophages (TAM) directly isolated from intestinal adenomas. This approach indeed unravelled a central role of *Hif1a* in TAMs for the expression of various fibroblast-activating factors, e.g. COX-2, IGF-1, IL-1β, IL-6 and granulin (Figure 5B). In our analyses, the expression of TGF-β1, the archetypical pro-fibrotic factor, was constantly lower in *Hif1a*-deficient TAMs, but did not reach statistical significance (Figure 5B). As the activation of TGF-β is a complex and highly regulated process (42), we hypothesized that HIF-1α is of importance in this setting. Indeed, HIF-1α -deficient AAMs displayed reduced levels of bioactive TGF-β in the supernatant (Figure 5C), suggesting a functional importance of HIF-1α for TGF-β activity elicited by TAMs. Next, we addressed the role of *Hif1a* for macrophage-mediated fibroblast proliferation (40, 43). While conditioned medium from AAM enhanced survival of primary murine intestinal fibroblasts (MIFs), no difference between WT and *Hif1a*-null macrophages was detectable (Figure S5B). Finally, we sought to address the secretion of pro-tumorigenic cytokines, another tumor-promoting function of TAFs (44). Stimulation of primary MIFs with conditioned medium from AAM induced gene expression of IL-6, HGF and epiregulin (*Ereg*), factors with established tumor-promoting activity in the intestine (45, 46). Of note, this effect was significantly reduced upon deletion of *Hif1a* in macrophages (Figure 5D).

**Figure 5.**
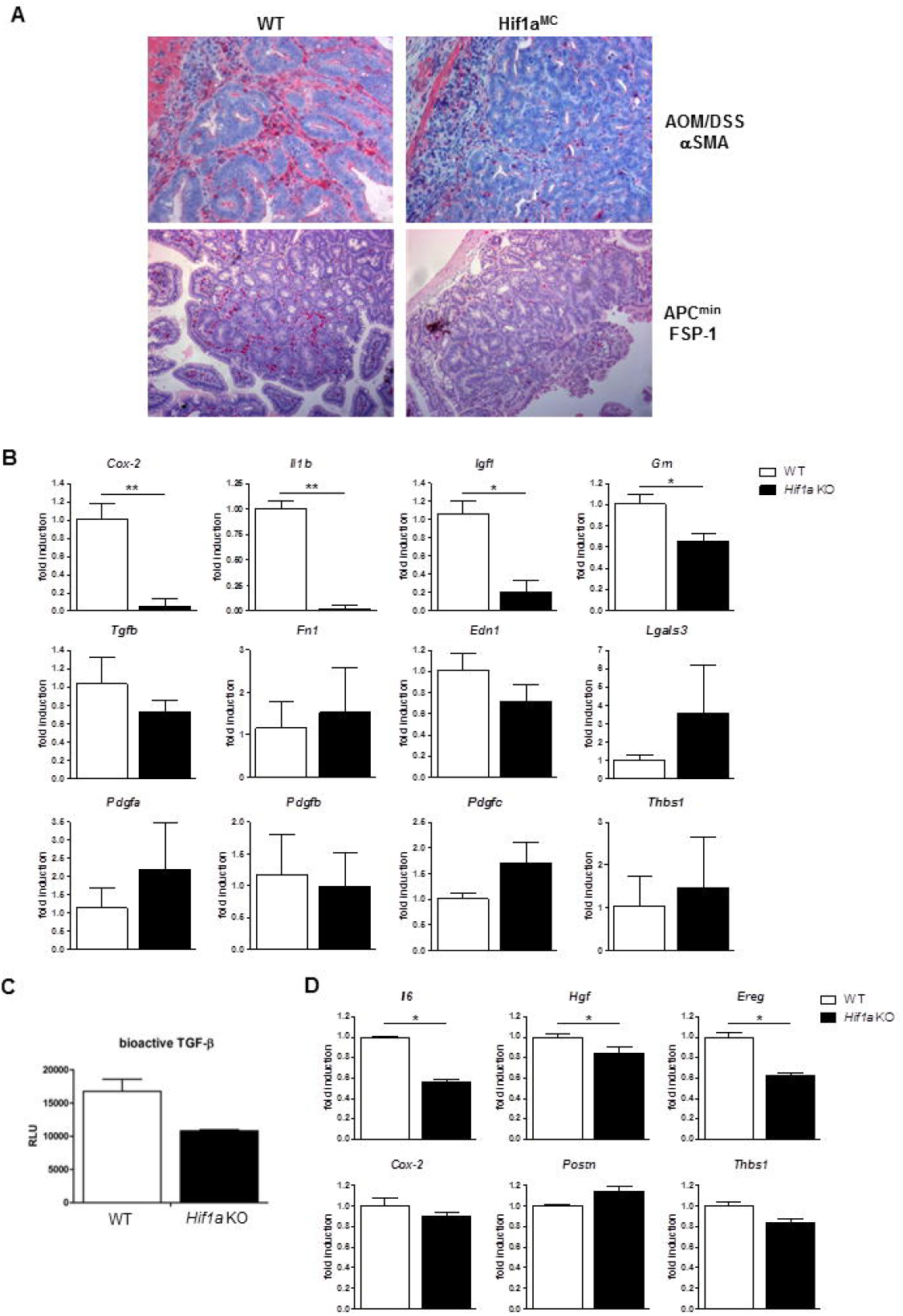
*Hif1a* in myeloid cells is essential for activation and pro-tumorigenic gene expression of intestinal fibroblasts. (**A**) Immunohistochemical analysis of myofibroblast markers aSMA and FSP-1 in intestinal sections from AOM/DSS-treated (above) and APC^min^ (below) WT and Hif1a^MC^ mice. (**B**) Relative mRNA expression of pro-fibrotic genes in tumor-associated macrophages isolated from APC^min^ adenomas from WT (*n* = 2) and Hif1a^MC^ (*n* = 4) mice. (**C**) Determination of bioactive TGF-β in the supernatant of WT and alternatively activated *Hif1a*-KO macrophages (*n* = 3 per group). (**D**) Relative mRNA expression of pro-tumorigenic genes in primary murine intestinal fibroblasts stimulated with conditioned media from WT and *Hif1a*-KO macrophages (*n* = 3 per group). Data are represented as mean + SEM. *p<0.05, **p<0.01 by unpaired Student’s t test.

### Myeloid-mediated activation of TAF precursor cells depends on *Hif1a*

TAFs can originate from different cellular sources, amongst others pericytes, mesenchymal stems cells (MSCs) and fibrocytes (44). We sought to investigate if macrophages are able to polarize these cell types into myofibroblasts and whether *Hif1a* is important in this setting. Gli1-positive pericytes, marked by tdTomato expression (47), as well as MSCs readily polarized into myofibroblasts after addition of macrophage conditioned medium (CM, Figure 6A and B). Intriguingly, HIF-1α was of central importance in this setting as the effect on pericytes was completely and that on MSCs partially abolished upon *Hif1a* deletion. Fibrocytes are of myeloid origin, display features of monocytes as well as fibroblasts and contribute to fibrosis in various organs and tumors (48, 49). We took advantage of an established protocol to generate fibrocytes *ex vivo* from splenic monocytes (50). Of note, *Hif1a*–deficient monocytes displayed greatly reduced capacity for fibrocyte production (Figure 6C). Furthermore, the expression of various tumor-supporting factors in fibrocytes was found to be regulated by *Hif1a* (Figure 6D). Taken together, these results point to a hitherto unknown function of *Hif1a* for the activation of TAF precursor cells of different origin.

**Figure 6.**
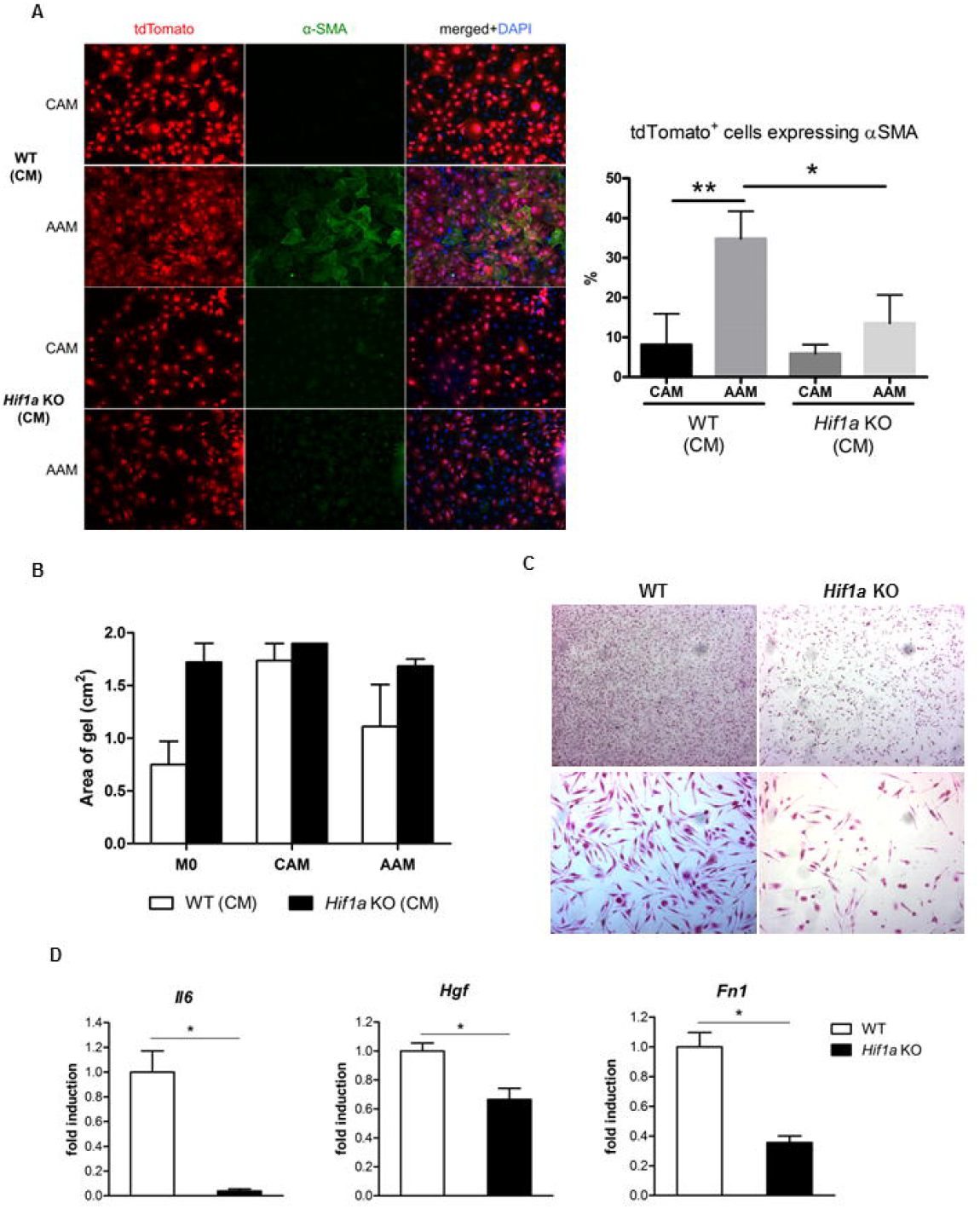
Myeloid cell-mediated activation of TAF precursor cells depends on *Hif1a*. (**A**) Representative images and quantification of Gli1^+^ MSC isolated from bone-chips of bigenic Gli1CreER^t2^;tdTomato mice and cultured with supernatants of CAM or AAM from either wildtype or Hif1a^MC^ mice and stained for alpha smooth muscle actin (αSMA) indicating myofibroblast differentiation (*n* = 3 per group). (**B**) Mesenchymal stem cells were cultured in collagen gels for 14 days either in stem cell expansion medium (SCEM, control) or in a mixture of SCEM and supernatants of M0, CAM or AAM (*n* = 3 per group). Collagen gel areas were measured using the ImageJ software. (**C**) Representative images of fibrocytes differentiated in vitro from splenic monocytes from WT and and Hif1a^MC^ mice. Magnification 25x (upper panel) and 100x (lower panel). (**D**) Relative mRNA expression of selected pro-tumorigenic factors in differentiated fibrocytes from WT and and Hif1a^MC^ mice (*n* = 3 per group). Data are represented as mean + SEM. *p<0.05 by unpaired Student’s t test.

### Non-canonical stabilization of HIF-1α predominates in murine intestinal tumors

The luminal cell layer of the colon is characterized by physiological hypoxia (51). Nuclear HIF-1α protein is readily detectable in luminal enterocytes (Figure S6A), suggesting hypoxia-induced HIF-1α protein stabilization in the gut under physiological conditions. Against this background, we sought to determine the relevance of hypoxia for HIF-1α protein stabilization in murine intestinal tumors. Intriguingly, hypoxic areas were detected only sporadically in AOM/DSS and APC^min^ adenomas (Figure 7A and B). This pattern of hypoxia was clearly not able to explain the pervasive stabilization of HIF-1α protein in both tumor types. Against this background it is relevant to note that hypoxia-independent means of HIF-1α protein stabilization have been identified and are gradually receiving more attention (25, 27). As activation of various oncogenes has been shown to result in hypoxia-independent HIF-1 stabilization, we decided to investigate the role of oncogenes and tumor suppressor genes characteristic for colon cancer (27, 52). To mimic the molecular events that instigate tumor formation in the APC^min^ model, mice with inducible *Apc* deletion were analyzed (53). Of note, acute *Apc* loss resulted in enhanced HIF-1α protein stability (Figure 7D). To analyze the role of oncogene activation for HIF-1α stabilization, we took advantage of transgenic mice with inducible, epithelial-specific expression of oncogenes with relevance for CRC pathogenesis (54). The strongest effect was noted for PIK3CA^H1047R^, the activation of which resulted in robust HIF-1α protein stabilization of the entire epithelial cell lining (Figure 7E). Activation of oncogenic KRAS or β-catenin resulted in localized HIF-1α stabilization in luminal epithelial cells (Figure 7F,G,A). These results point towards a functional role of key tumor suppressors and oncogenes regulating the Wnt/β-catenin, PI3K and MAPK cascades for non-canonical HIF-1α stabilization during intestinal tumor formation and progression.

**Figure 7.**
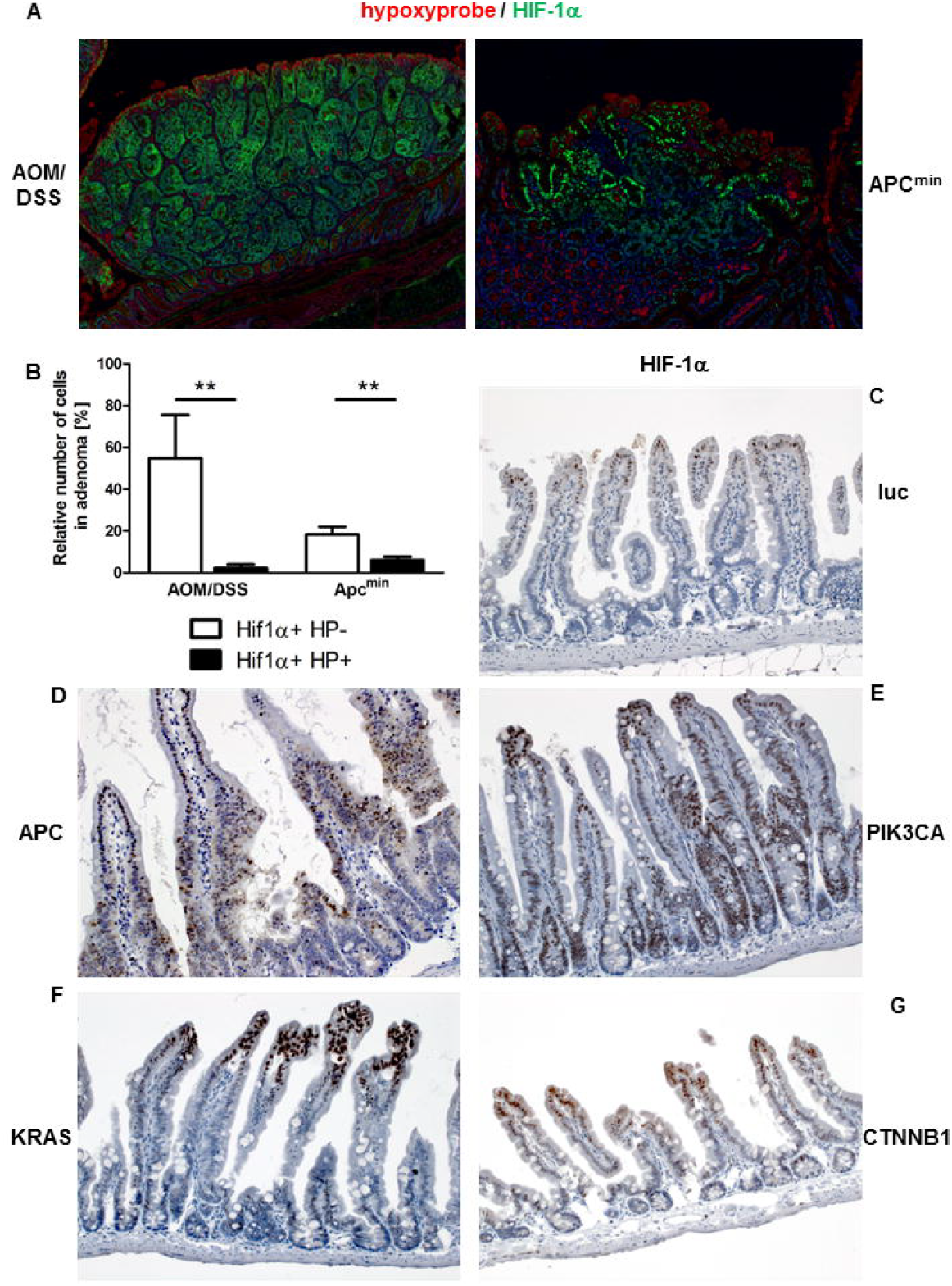
Non-canonical stabilization of HIF-1α in murine intestine. (**A**) Immunofluorescent analysis of hypoxyprobe (red) and HIF-1α (green) in AOM/DSS-induced (left) and APC^min^ (right) adenomas. (**B**) Analysis of relative number of cells in adenomas expressing either hypoxyprobe (HP) or HIF-1α. (**C**-**G**) Immunohistochemical staining of HIF-1α in intestinal sections from mice with inducible expression of firefly luciferase (**C**, luc), inducible loss of APC (**D**), PIK3CA (**E**), KRAS (**F**) or oncogenic β-catenin (CTNNB1, **G**). Data in B show mean + SD, 7-8 adenomas from 2 animals per model were evaluated. **p<0.001 by unpaired Student’s t test.

## Discussion

Here, we addressed the functional importance of HIF-1 for colon cancer in a cell type-specific manner, using transgenic mice harbouring either an intestinal epithelial cell-(Hif1a^IEC^) or a myeloid cell-specific (Hif1a^MC^) *Hif1a* deletion (21, 22, 26). We found that *Hif1a* plays multiple non-redundant roles in these two key cell types of intestinal tumors. Our finding of reduced tumor growth in Hif1a^IEC^ mice is in line with earlier reports showing *Hif1a*-dependent growth of human CRC xenografts (11, 12). Also in line with our results, the group of Celeste Simon reported that application of a chemical HIF inhibitor decreased tumor formation in mice bearing AOM/DSS adenomas and s.c. CT26 allografts (13). Of note, transgenic overexpression of oxygen-stable HIF-1α in intestinal epithelial cells did not further accelerate the formation of AOM/DSS and APC^min^ tumors (14). Deletion of *Hif1a* in IECs did not affect tumor size in a chemical model of proximal colon cancer (16), arguing for the importance of cell type- and location-specific factors that need to be further investigated.

In our experiments, Hif1a^IEC^ mice displayed lower levels of pro-inflammatory cytokines in acute DSS-induced colitis, suggesting an activating function of HIF-1 in IECs for intestinal inflammation. This result contrasts with earlier reports from various independent groups. Karhausen *et al.* noted enhanced activity in hapten-induced intestinal inflammation upon conditional loss of *Hif1a* in IECs (51) and identified reduced barrier integrity in mutant mice as the underlying mechanism. Against this background, we determined intestinal barrier function in our mice and could not find a difference between WT and Hif1a^IEC^ mice (Figure S1B). Later, Shah *et al.* reported no effect of IEC-specific *Hif1a* loss on the severity of acute DSS-induced colitis (55). It is well established that the susceptibility to experimentally induced colitis is genetically determined and differs substantially between inbred strains of mice (56). Karhausen *et al.* and Shah *et al.* used mice with a mixed genetic background while our mice were >99% C57Bl6/J, potentially explaining the different results. Additional support for a protective role of HIF-1 during intestinal inflammation came from two simultaneously published seminal reports by the groups of Sean Colgan and Cormac Taylor showing that inhibitors of prolyl hydroxylases (PHDs), a group of enzymes crucial for HIF degradation, protect against murine colitis (57, 58). In our opinion, different explanations are possible regarding the conflict with our data, e.g. a functional importance of other PHD targets, e.g. NF-κB, and hydroxylase-independent functions of PHD inhibitors. Furthermore, systemically administered PHD inhibitors target every cell they encounter while in our genetic model HIF-1 was exclusively deleted in IECs, precluding a direct comparison of the different experimental approaches.

Our data support a central role of HIF-1α in myeloid cells for the activation of fibroblasts in the stroma of intestinal tumors. In the context of wound healing and organ fibrosis, the importance of macrophages, especially that of alternatively activated macrophages (AAM), for fibroblast activation is well established (59, 60). AAMs are of special significance as they express various fibroblast-activating factors (40). While earlier reports showed a functional relevance of HIF-1 for the control of gene expression of pro-fibrotic factors (e.g. TGF-β1, endothelin-1, fibronectin-1 and COX-2 (61-64)) in different cell types, *Hif1a* deletion in AAMs remained without effect on mRNA expression of a comprehensive set of pro-fibrotic factors in our experimental setup. To address the importance of the tissue context, we isolated tumor-associated macrophages (TAMs) from intestinal adenomas. Of note, in these cells the expression of various pivotal pro-fibrotic genes was indeed regulated by *Hif1a*. These results not only underscore the significance of *Hif1a* for TAM-mediated activation of TAFs, but again illustrate the importance of experimental conditions that more closely resemble the tissue context.

TAFs were for the longest time unequivocally considered to support tumor growth and progression (44). This concept was challenged by a report from the group of Raghu Kalluri, showing accelerated tumor progression upon genetic ablation of α smooth muscle actin-positive TAFs in murine pancreatic ductal adenocarcinoma (65). At first glance, this observation contrasts with our data arguing for a tumor-supporting role of TAFs. Besides cancer type- and context-specific factors, the different experimental approaches are strong candidates for potential explanations. The Kalluri group used an elegantly designed genetic model resulting in direct ablation of TAFs. Our experimental setup, on the other hand, targeted TAFs indirectly via deletion of *Hif1a* in myeloid cells. The study by Ozdemir *et al.* points towards previously unappreciated tumor-inhibiting functions of TAFs. How pro- and anti-tumor aspects of TAFs are regulated is an intriguing question. While our data suggest that myeloid *Hif1a* impacts mainly on tumor-supporting aspects of TAFs, future work has to address the eligibility of HIF-1 inhibitors in the setting of a stroma-targeting cancer therapy.

Our study highlights that HIF-1α can be stabilized by multiple means in the colon cancer context. We show that loss of APC, as well as oncogenic activation of PIK3CA or KRAS can contribute to HIF-1α stabilization, even in the absence of overt hypoxia. Our findings give credence to the concept of non-canonical HIF-1α stabilization that was recently coined by Amato Giaccia and colleagues (25). We have not determined the exact molecular link(s) relaying oncogenic events to HIF-1α stabilization in this study. However, our data add a further layer of complexity to HIF-1α regulation, and suggest that hypoxia and HIF-1α stabilization can be uncoupled in cancer.

## Methods

### Animals and Mouse models

All mice were housed in a specific pathogen-free environment and were allowed to adapt for at least 7 days to housing conditions before starting an experiment. Mice were given tap water and standard rodent chow *ad libitum* and were housed at constant room temperature with a 12 hour light/dark cycle. All experiments were performed by using weight-, age- and sex-matched littermates (VillinCre/*Hif1a*^loxP/loxP^, VillinCre/*Hif1a*^*l*oxP/loxP^/APC^+/min^, LysMCre/*Hif1a*^loxP/loxP^ and LysMCre/*Hif1a*^loxP/loxP^/APC^+/min^, all on a C57BL/6J background). For the AOM/DSS model, 6-8 week-old mice were injected intraperitoneally with 10 mg/kg AOM (azoxymethane; Sigma-Aldrich, Germany) followed by three cycles of 2% dextran sodium sulfate (DSS; MP Biomedicals, Germany) in drinking water for five days and normal drinking water for 16 days. Mice were sacrificed 8 weeks after AOM injection. Tumors were counted and measured under a dissecting microscope and colonic normal and tumor tissue snap frozen in liquid nitrogen for *ex vivo* analysis. In the acute DSS model, 10-12 week-old mice received 2% DSS in the drinking water for 6 days and were sacrificed on the last day of DSS administration. In order to assess DSS colitis activity, body weight, stool consistency and the presence of occult or gross blood were determined and a scoring system was applied (66). For tumor counting in the APC^min^ model, mice were sacrificed at 80 days of age. The entire intestine was removed and the small intestine divided into three equal sections, with a fourth section comprising the entire large intestine. Intestinal segments were opened longitudinally and fixed at 4°C for 36 h in 10% neutral buffered formalin. Adenomas were visualized by staining with 0.1% methylene blue and counted and measured under a dissecting microscope. For histology, small intestines and colons were removed, flushed with PBS, fixed in 10% neutral buffered formalin at 4°C overnight and paraffin-embedded.

Mice with inducible oncogenes, transgene cassettes contain doxycycline-inducible tdTomato linked via 2A peptides to human PIK3CA(H1047R), KRAS(G12V) or firefly luciferase, and a PGK promoter flanked by heterologous loxP and lox5171 sites. Transgenes were integrated by Cre recombinase-mediated cassette exchange into a previously modified Gt(ROSA)26Sor locus harboring a promoterless neomycin resistance gene in F1 hybrid B6/129S6 embryonic stem cells. Recombined clones were selected by 250 μg/ml G418 and analysed by Southern blot of genomic DNA digested with BamH1 and HindIII, using the neomycin cassette as a probe (67). Animals were generated by diploid aggregation. Transgenes were induced *in vivo* by administration of doxycycline (4 mg/ml) provided in a 5% sucrose solution via the drinking water. Mouse lines were inbred to C57Bl6/J.

### FDG PET/CT

PET (positron emission tomography) was performed longitudinally at 6 weeks after initiation of AOM/DSS treatment in 14 mice (7 WT and 7 Hif1a^IEC^) after i.v. injection of 27.1 ± 7.4 MBq F-18-flurodeoxyglucose (FDG). Region of interest analysis was performed by manual lesion segmentation in the PET data co-registered to a whole-body CT (computational tomography) using the PMOD 3.4 software package (Technologies Ltd, Zürich, Switzerland). The maximum FDG uptake in the lesion was scaled to the mean FDG uptake in the thigh muscles.

### Metabolomics extraction

Colon samples were homogenized in two cycles with a 20 s long cooling phase on ice in between by Precellyse24 (Bertin Technologies, France). Volumes of the ice-cold extraction solution methanol-chloroform-water (all Sigma, 5:2:1 v/v/v) were adjusted to the sample weight (1 ml/50 mg tissue). Extraction solvent contained an internal extraction standard cinnamic acid (Sigma, 2 μg/ml). Colon extracts were incubated for 30 min at 4°C on a rotary wheel, and subsequently centrifuged for phase separation (15 min, maximum speed). Equal volumes of polar phase were selected from each sample separately and dried under vacuum. Extracts were stored at -25°C until continuing with the preparation for GC-MS measurement.

### GC-MS sample preparation

Derivatization was carried out as described with modifications (68). At first dried extracts were dissolved in 20 μl of methoxyamine hydrochloride solution (Sigma, 40 mg/ml in pyridine (Roth)) and incubated for 90 min at 30°C under constant shaking. In a second phase, samples were incubated with 80 μl of N-methyl-N-[trimethylsilyl]trifluoroacetamide (MSTFA; Macherey-Nagel, Düren, Germany) at 37°C for 45 min. The in-house alkane standards were added prior to the MSTFA for retention time analysis (10 μl/ml MSTFA). The extracts were centrifuged for 10 min at 10,000g, aliquoted and 30 μl transferred into glass vials for GC-MS measurements. Formulation of standards for retention index determination, as well as for identification and quantification of metabolites are described in (69). Identification and quantification standards were prepared in parallel with samples and in the same manner.

### GC-MS measurement

Metabolite analysis was performed on a gas chromatography coupled to time of flight mass spectrometer (Pegasus III-TOF-MS-System, LECO Corp., St. Joseph, MI, USA), complemented with an auto-sampler (MultiPurpose Sampler 2 XL, Gerstel, Mülheim a.d. Ruhr, Germany). The samples and quantification standards were injected in split mode (split 1:5, injection volume 1 μl) in a temperature-controlled injector with a baffled glass liner (CAS4, Gerstel). The following temperature program was applied during sample injection: Initial temperature of 80°C for 30s followed by a ramp with 12°C/min to 120°C and a second ramp with 7°C/min to 300°C and final hold for 2 min. Gas chromatographic separation was performed on an Agilent 6890 N (Agilent, Santa Clara, CA, USA), equipped with a VF-5 ms column of 30-m length, 250-μm inner diameter, and 0.25-μm film thickness (Varian, Palo Alto, CA, USA). Helium was applied as carrier gas with a flow rate of 1.2 ml/min. Gas chromatography was performed with the temperature gradient: 2 min heating at 70°C, first temperature gradient with 5°C/min up to 120°C and hold for 30 s; subsequently, a second temperature step of 7°C/min up to 350°C with a hold time of 2 min. The spectra were recorded in a mass range of 60 to 600 U with 20 spectra/s at a detector voltage of 1.750 V.

### GC-MS data analysis

File processing and analysis was performed with the vendor software ChromaTOF Version 4.42 (LECO). Processing parameters: baseline offset of 1, peak width of 4 s, signal/ noise of 20, and peak smoothing of 11 data points. Retention indices were calculated based on retention index standards. The Golm Metabolome Database (GMD) provided mass spectra and retention information for peak identification (70). Quantification of metabolites was done by external calibration as described in (69).

### Differentiation of murine fibrocytes

Murine fibrocytes were cultured and differentiated as described in detail before (50). Briefly, mouse splenocytes were isolated by forcing spleens through a 100 μm cell strainer (BD Biosciences, Heidelberg, Germany), collected by centrifugation (300 x g for 5 min) and further purified by erythrocyte lysis. After centrifugation, CD11b-positive monocytes were enriched from spleen cells using CD11b MicroBeads (Miltenyi Biotec, Bergisch Gladbach, Germany) according to the manufacturer’s protocol. Monocyte-enriched cells were resuspended in Fibrolife basal media (Lifeline Cell Technology, Frederick, USA) supplemented with 10 mM HEPES (Gibco), 2 x non-essential amino acids (Biochrom), 2 mM sodium pyruvate (Biochrom), 4 mM glutamine (Gibco), 100 U/ml penicillin, 100 g/ml streptomycin, 2× ITS-3 (Sigma-Aldrich) and 50 μM 2-mercaptoethanol (Gibco). For fibrocyte differentiation, cells were cultured in the presence of 50 ng/ml murine IL-13 (eBioscience) and 25 ng/ml murine M-CSF (eBioscience). On day 3 of the incubation, wells were further supplemented with a cocktail containing a final concentration of 25 ng/ml IL-13 and 12.5 ng/ml M-CSF in fibrolife media. 5 days later, cells were fixed with 10% neutral buffered formalin and stained with hematoxylin and eosin or cell pellets were snap frozen for RNA extraction.

### Isolation and culture of mouse Gli1+ cells

Gli1CreERt2 (i.e. Gli1tm3(re/ERT2)Alj/J, JAX Stock #007913), Rosa26tdTomato (i.e. B6-Cg-Gt(ROSA)26Sorttm(CAG-tdTomato)Hze/J, JAX Stock #007909) were purchased from Jackson Laboratories (Bar Harbor, ME). Bigenic Gli1CreERt2;tdTomato at 8 weeks of age received tamoxifen (3× 10mg) in corn oil / 3% ethanol (Sigma) via oral gavage and were sacrificed 10 days after the last tamoxifen dose. Gli1+ bone marrow mesenchymal stem cells (MSC) were isolated from compact bone chips of femur, tibia and pelvis according to the isolation protocol by Zhu *et al.* (71). Gli1+ cells were grown in alpha MEM (GlutaMAX, Life Technologies) containing 20% MSC qualified FBS (Life Technologies), 2% Penicillin Streptomycin (Life Technologies), 1ng/ml murine basic fibroblast growth factor (Thermo Fisher Scientific) and 5ng/ml murine epidermal growth factor (Peprotech). FACS sorting was performed using the FACSAria II cell sorter (BD Biosciences) in passage 2. To study the effect of macrophage supernatant on myofibroblast differentiation of Gli1+ cells, cells were grown on coverslips (Fisher), serum starved overnight (0.5% FBS) and treated with pure supernatant of classically or alternatively activated macrophages from wildtype or Hif1a^MC^ mice for 48 hours. Cells were stained with a FITC conjugated mouse anti α-SMA antibody (1:200, Sigma, # F3777). The number of positive cells per high power field was quantified using a Keyence BZ9000 fluorescence microscope.

### Murine intestinal fibroblasts

Intestinal fibroblasts were isolated from murine intestine as described in detail before (72). Briefly, intestine was detached from the mesenterium and epithelial cells were denuded by washing twice in 5mM EDTA/HBSS for 15 min followed by dispase and collagenase D incubation for another 30 min. Epithelial cell-denuded mucosal samples were resuspended in DMEM (Gibco) supplemented with 5% FBS, and cultured under 5% CO_2_. After one week, passaged fibroblasts survived whereas epithelial cells and macrophages underwent senescence. Cells were characterized by positive immunostaining of α-smooth muscle actin. Intestinal fibroblasts of passage 2-5 were used for all experiments. For RNA isolation, 150.000 cells/well were seeded in 6-well plates and cultured in DMEM supplemented with 1% FBS overnight. On the next day, cells were stimulated with macrophage-conditioned media. 24 h later, RNA isolation was performed using TRIzol (Invitrogen) according to the manufacturer’s instructions.

### Culture and stimulation of small intestinal organoids and APC^min^ spheroids

Based on protocols established by the Clevers group (73), a modified protocol (74) was used for crypt isolation from mouse small intestine. For organoid culture, the complete small intestine from 6-12 weeks old mice was used. Spheroid culture were grown from the distal third of the intestine from 15-20 weeks old APC^min^ mice. Villi were scraped off the tissue using coverslips. Tissue pieces were washed in PBS and incubated shaking in 1 mM EDTA/PBS on ice for 30 min and for one further hour in 5 mM EDTA/PBS. Subsequently, the crypt suspension was passed through a 70 μm cell strainer. Following centrifugation, crypts were resuspended in Matrigel (Corning). Polymerized Matrigel drops were overlaid with ENR (Egf/noggin/R-spondin) medium. Spheroids were cultured in medium without R-spondin, containing 10 μM ROCK inhibitor (Sigma). Medium was changed after 2-3 days and spheroids were no longer treated with ROCK inhibitor from this time point on. After 5-7 days incubation, organoids or spheroids were disrupted and re-seeded in Matrigel. Two days later, organoids were used for experiments, in which they were treated with 450 ng/ml LPS or 100 ng/ml murine TNFα and exposed to hypoxia (1% O_2_) for 6 hours before RNA isolation. Spheroids were treated with 1:5 diluted BMDM supernatants directly after passaging and monitored for up to five days. Spheroid diameter and number were calculated by evaluation of microscopic pictures using ImageJ software.

### Isolation of F4/80+ cells from intestinal adenomas

Small intestinal adenomas were isolated from 120 days old APC^min^ mice. The tissue was cut into small pieces and digested with Collagenase C and DNase. For further disruption, it was pulled through 18G and 22G needles. Cells were collected through a 70 μm cell strainer and incubated in serum-containing buffer to avoid unspecific antibody binding. In the next step, cells were labeled with biotinylated anti-F4/80 antibody (BioLegend) before magnetic Streptavidin MicroBeads (Miltenyi, Bergisch Gladbach, Germany) were added. Thereafter, F4/80-positive cells were selected by magnetic separation using LS columns (Miltenyi, Bergisch Gladbach, Germany) and finally dissolved in PeqGold RNA Pure reagent (Peqlab, Erlangen, Germany) for RNA isolation.

### RNA extraction and quantitative PCR analysis

Intestinal tissue was disrupted and pulverized with a TissueLyser II (Qiagen, Hilden, Germany). Total RNA was isolated using the RNeasy Mini Kit (Qiagen) and QIAshredder spin columns (Qiagen) according to the manufacturer’s instructions. The yield and purity of RNA were determined spectrophotometrically. RNA integrity was checked by MOPS buffered denaturizing RNA gel electrophoresis and only samples with a 28S/18S ratio of at least 1.8 were included in the qPCR study. First strand cDNA of 1 or 2 μg total RNA/per reaction was synthesized with an oligo (dT) primer and a SuperScript™ First Strand Synthesis System (Invitrogen) according to the manufacturer’s recommendations. Quantitative real-time PCR analysis was conducted on an ABI 7500 Real-Time PCR system using PowerSYBR^®^ Green PCR Master Mix (Thermo Fisher Scientific, Waltham, MA USA). Primer-specific annealing temperatures were pre-evaluated prior to the study to optimize PCR conditions. Primer specificity was checked by melt-curve analyses and TAE-buffered DNA agarose gel electrophoresis of obtained PCR products. Amplification efficiency was calculated with LinRegPCR 2016.0 (Heart Failure Research Center, Amsterdam, The Netherlands) and used for efficiency corrections (75). Inter-Run calibrators were used if necessary to correct for inter-run variations. Relative fold-changes of target gene expression were calculated with qbase+ 3.0 (Biogazelle, Zwijnaarde, Belgium) that automatically applies the above mentioned corrections to the ΔΔCq method (76). Primers are listed in the supplemental information.

### Immunohistochemistry

Immunohistochemical analysis was conducted on formalin-fixed paraffin-embedded tissues. For histopathological evaluation, intestinal sections were stained with hematoxylin and eosin and mucosal inflammation was graded as described (66). Intestinal sections were also stained with antibodies specific for F4/80 (Bio-Rad, Hemel Hempstead, UK), FSP-1 (Dako, Hamburg, Germany), HIF-1α (Cayman Chemical, Ann Arbor, USA), αSMA (Merck Millipore, Darmstadt, Germany) and ZO-1 (Thermo Fisher Scientific, Waltham, USA). The antibody-antigen complexes were detected using biotinylated donkey anti-rat and donkey anti-rabbit secondary antibodies (Dianova, Hamburg, Germany) and the Dako REAL Detection System (Dako). Sections were counterstained with hematoxylin. Immunohistochemical detection of HIF-1α was performed as previously described (77). For the immunohistochemical assessment of hypoxic areas, the Hypoxyprobe-1 Kit (Hypoxyprobe Inc., Burlington, USA) was used as described before (78). Tumor-associated macrophage abundance was determined by quantification of F4/80+ cells in at least 5 random fields of view.

### Multiplex immunohistochemistry

Paraffin sections were deparaffinized and rehydrated according to standard protocols. Subsequently, the sections were fixed by 10 min incubation in 3.5% formalin. Antigen retrieval was achieved by 15 min cooking at 110°C in Dako target retrieval solution using the Decloaking Chamber (Biocare Medical). After 10 min blocking in Dako antibody diluent, the sections were incubated with anti-HIF-1α (Cayman Chemical, 1:20.000 in Dako antibody diluent) for 30 min and afterwards treated with ImmPRESS anti-rabbit IgG (Vector) for 20 min. To permanently label Hif1α with a fluorescent tag, sections were treated with Opal 520 (Opal 4-color IHC kit, PerkinElmer, 1:50 in provided amplification reagent) for 10 min. Subsequently, the antibodies were removed from sections by microwave treatment in AR6 buffer (Opal 4-color IHC kit, PerkinElmer), while the bound Opal fluorophore outlasted this procedure. For additional staining of Hypoxyprobe (pimonidazole adducts), the sections were incubated with mouse IgG block (M.O.M. detection kit, Vector) over night at 4°C. At the next day, anti-pimonidazole mouse IgG (hpi, 1:200 in 1:10-diluted protein concentrate from M.O.M. kit) was added to the sections for 30 min. Thereafter, sections were first incubated with ImmPRESS mouse IgG (Vector) for 20 min and then treated with Opal 670 (Opal 4-color IHC kit, PerkinElmer, 1:50 in provided amplification reagent) for 10 min. After repeated microwave treatment, nuclei were stained with spectral DAPI (Opal 4-color IHC kit, PerkinElmer) for 5 min. Coverslips were mounted on the stained tissue sections using Vectashield HardSet mounting medium (Vector). Fluorescent signals were detected, separated and recorded using the Vectra 3.0 multiplex imaging system (PerkinElmer).

### Isolation and stimulation of bone marrow-derived macrophages (BMDM)

Bone marrow was collected from tibiae, femurs and humeri of 8-12 weeks old wildtype and Hif1a^MC^ mice. After flushing out the marrow, red blood cells were lysed with ACK buffer and cells were seeded on plastic plates in RPMI supplemented with 10% FBS, 100 U/ml penicillin, 100 μg/ml streptomycin. Next day, non-attached cells were collected and cultured in RPMI supplemented with 20% FBS and 30% L929-conditioned medium for one week. Differentiated BMDMs were stimulated for 48 h with LPS (100 ng/ml, Sigma Aldrich) and γ-IFN (20 ng/ml) for classically activated (CAM) and with IL-4 (20 ng/ml, both from eBioscience) for alternatively activated macrophages (AAM). For RNA isolation, cells were harvested using TRIzol (Invitrogen) from M0 (non-polarized), CAM and AAM. To collect conditioned media from polarized macrophages, cells were extensively washed with PBS 48 h after stimulation and fresh media was added for an additional 24 h.

### Flow cytometric analyses of murine leukocyte populations

Small intestine was rinsed thoroughly and incubated in buffer containing EDTA to remove the mucus. Cells were minced and digested with type IV collagenase (Cellsystems). Blood was taken from the right ventricle. Small intestine, blood, bone marrow, and spleen cells were subjected to red blood cell lysis using Pharm Lyse (BD Biosciences, San Jose, USA). Flow cytometric analysis was done as described in detail before (79). Antibodies were purchased from eBioscience (CD45, CD11b, Ly6G, F4/80, CD3, CD4, CD8, CD103, and NK1.1), BD Biosciences (Gr1, I-A^b^), and Biolegend (CD11c).

### Statistics

Statistical analysis was done using Prism 4.0 software (GraphPad Software, San Diego, California, USA). Statistical significance was determined by two-tailed Student’s *t* test for unpaired or paired observations. Differences were considered statistically significant at *p*<0.05. Data are expressed as means + SEM. The asterisks in the graphs indicate statistically significant changes with *p* values: * *p* < 0.05, ** *p*≤0.01 and *** *p*≤0.001.

### Study approval

Experimentation and transgenic animal generation was approved by authorities in Berlin (Landesamt für Gesundheit und Soziales: G0004/07, G0185/09, G0143/14).

## Author Contributions

Conceptualization, N.R., M.M. and T.C.; Methodology, N.R., M.E., S.J., A.E., K.T.W., A.A.K., A.F., S.N., M.G., I.R., T.E., C.Z., S.K., R.H., M.B.M., W.F. and M.M.; Formal Analysis, Investigation, and Visualization, N.R., M.E., S.J., A.E., A.A.K., R.K., S.N., I.R., M.G., C.Z., S.K., M.B.M., M.M. and T.C.; Writing, N.R., F.T., M.M. and T.C.; Supervision, Resources, and Funding Acquisition, R.K., S.K., C.E.L., W.B., M.S., L.S., O.S., F.T., M.M. and T.C.; Project Administration, T.C.

## Acknowledgements

Research in the Cramer lab was supported by grants from Deutsche Krebshilfe (109160) and Deutsche Forschungsgemeinschaft (CR 133/2-1 until 2-4). Nadine Rohwer was supported by a grant from the BMBF (MAPTor-NET (031A426A)). We are indebted to Deborah Gumucio (University of Michigan, USA) for providing Villin-Cre mice. We are grateful to Christine Sers (Charité, Berlin) and Florian R. Greten (Georg-Speyer-Haus, Frankfurt) for helpful discussions and to Ilia N. Buhtoiarov (Children’s Hospital, Cleveland, USA), Glenn S. Belinsky, Daniel W. Rosenberg (University of Connecticut Health Center, Farmington, USA) and Takuji Tanaka (Gifu Municipal Hospital, Japan) for providing control reagents. We very much appreciate the help of Ralf Weiskirchen (RWTH University Hospital Aachen) with the MLEC assay. The excellent technical assistance of Birgit Bogdanoff and Simone Spiekermann is highly appreciated.

